# Anticodon-edited tRNA enables translational readthrough of *COL4A5* premature termination codons

**DOI:** 10.1101/2025.08.07.669229

**Authors:** Kohei Omachi, Joseph J. Porter, John D. Lueck, Jeffrey H. Miner

## Abstract

Alport syndrome is caused by variants in *COL4A3*, *COL4A4*, or *COL4A5*, which encode the α3α4α5 chains of type IV collagen. These variants result in defects in the glomerular basement membrane (GBM) and impaired kidney function. Nonsense variants result in truncated proteins lacking the NC1 domain, thereby preventing proper GBM assembly and function and causing the most severe forms of the disease. Restoring full-length protein expression represents a potential therapeutic strategy for Alport syndrome and related disorders. Anticodon-edited transfer RNAs (ACE-tRNAs), which promote premature termination codon (PTC) readthrough, have shown promise in diseases such as cystic fibrosis, but their application in Alport syndrome remains unexplored. To assess the potential of ACE-tRNAs for PTC readthrough of *COL4A5* nonsense variants, we employed a C-terminal NanoLuc-fused *COL4A5* reporter system in which luminescence is produced only upon full-length protein translation. We introduced ACE-tRNAs into HeLa and 293T cells expressing one of four *COL4A5* nonsense variants (*S36X, R1563X, S1632X, and R1683X*) identified in patients with X-linked Alport syndrome. Readthrough efficiency was evaluated via NanoLuc luminescence and western blotting. Furthermore, we assessed the efficiency of ACE-tRNA-restored α3α4α5 heterotrimer formation using a split NanoLuc-based assay. Our results show that application of ACE-tRNAs led to restored C-terminal luminescence across all four COL4A5 nonsense variants, indicating successful readthrough and full-length translation. Moreover, the restored COL4A5 proteins formed α3α4α5 heterotrimers. These findings support ACE-tRNA-mediated nonsense suppression as a promising therapeutic strategy for Alport syndrome, with the potential to restore GBM integrity in patients harboring nonsense variants.

## Introduction

Nonsense variants, which introduce premature termination codons (PTCs), result in the production of truncated proteins that are often nonfunctional and are therefore frequently associated with severe phenotypes across a wide range of genetic diseases, including cystic fibrosis [1] and Duchenne muscular dystrophy [2]. One promising therapeutic approach is PTC readthrough, in which the translational machinery bypasses the premature termination codon, allowing for the synthesis of full-length proteins. This phenomenon has been extensively investigated, particularly in the context of small molecules that can induce readthrough. Among these, aminoglycosides such as gentamicin have long been recognized as PTC readthrough inducers [3–5]. In recent years, derivatives of aminoglycosides with improved readthrough efficiency and reduced toxicity have been developed, offering potential clinical benefits [6, 7]. Despite these advancements, achieving therapeutically meaningful levels of readthrough remains a major challenge. To address this limitation, combination approaches incorporating readthrough potentiators have emerged as a complementary strategy [8, 9]. These compounds enhance the activity of primary readthrough agents and may enable lower, less toxic dosages while improving functional protein restoration.

Another emerging strategy to bypass premature termination codons involves the use of anticodon-edited transfer RNAs (ACE-tRNAs)[10–12]. These engineered tRNAs carry a modified anticodon that enables them to specifically recognize premature stop codons and insert the original amino acid that should be present in the wild-type protein, thereby restoring the synthesis of full-length, functional proteins. Compared to aminoglycosides and their derivatives, ACE-tRNAs exhibit significantly higher readthrough efficiency [13]. Aminoglycoside-induced readthrough depends on near-cognate tRNAs, which often insert amino acids that differ from the original [14], and are active at both sense and nonsense codons [15, 16]; thus, the resulting proteins may contain substitutions that impair function or stability. In contrast, ACE-tRNAs offer greater fidelity by directing the insertion of the correct amino acid, enhancing the likelihood of producing fully functional proteins.

Alport syndrome is the second most common genetic kidney disease. It is characterized by a glomerular filtration defect followed by progressive chronic kidney disease that eventually leads to kidney failure [17, 18]. Alport syndrome is caused by loss-of-function variants in one of the genes encoding the type IV collagen α3α4α5 network—*COL4A3* [19, 20], *COL4A4* [21], or *COL4A5* [22]. Among the many variants found in these genes, nonsense variants represent approximately 6% of cases. While missense variants are more common, nonsense variants are typically associated with a more severe clinical phenotype due to their impact on protein truncation and loss of function.

The type IV collagen α3α4α5 is synthesized as a heterotrimer, initiated through interactions at the C-terminal non-collagenous (NC1) domain, followed by stabilization via the collagenous domain and the N-terminal 7S domain [18]. In the case of nonsense variants, the resulting premature termination codon leads to truncated collagen chains that lack the NC1 domain, making them incapable of trimerization and thus unable to form a functional GBM scaffold. Given this pathogenic mechanism, therapeutic strategies aimed at restoring full-length type IV collagen proteins provide a rational and potentially disease-modifying treatment avenue for individuals with nonsense variant-associated Alport syndrome. Although ACE-tRNA therapy has been explored in various diseases using cultured cells [10, 13, 23–28] and mouse models [29, 30], its potential application in Alport syndrome has not yet been investigated.

In this study, we evaluated the therapeutic potential of ACE-tRNAs to promote readthrough of PTCs in *COL4A5*. Utilizing a NanoLuc-based translational reporter system, we demonstrated that ACE-tRNAs effectively induce full-length COL4A5 protein expression in cultured cells expressing nonsense mutation-containing cDNAs. Importantly, the restored full-length COL4A5 protein was capable of forming heterotrimeric complexes with COL4A3 and COL4A4, suggesting that the translation products are not only structurally complete but also functionally competent. These findings establish a proof-of-concept for ACE-tRNA-mediated therapy in nonsense variant-driven Alport syndrome, supporting further development toward in vivo therapeutic applications.

## Materials and Methods

### Cell culture

HeLa cells (ATCC, #CCL-2) were purchased from ATCC and maintained in Minimum Essential Medium (Gibco, # 11095080) supplemented with 10% FBS (Gibco, # A5670701), 1% penicillin streptomycin (Gibco, # 15140122) at 37 °C in a humidified 5% CO_2_ incubator. 293T cells were purchased from ATCC and maintained in Dulbecco’s Modified Eagle Medium (Gibco, # 10644633) supplemented with 10% FBS (Gibco, # A5670701), 1% penicillin streptomycin (Gibco, # 15140122) at 37 °C in a humidified 5% CO_2_ incubator. One day before transfection, cells were trypsinized and seeded into 6-well tissue culture plates (TPP, #92006). At 70-80% confluency, cells were transfected using Xfect transfection reagent (Takara, #631317) for HeLa and FuGENE 6 transfection reagent (Promega, # E2691) for 293T cells according to the manufacturer’s instructions. One day after transfection, the cells were trypsinized and seeded into Nunc 96-well white-bottom plates (Thermo Scientific, # 136102). Two days after transfection, cells were used for luciferase assays. For nonsense-mediated mRNA decay (NMD) inhibition experiments, cells were treated with 0.5 µM CC-90009 (Selleck, #S9832) for 24 hours. Luciferase assays were conducted using DMEM without phenol red (Gibco, #11054020).

### Plasmid DNA

Plasmids for the NanoLuc-based COL4A5 translational reporter assay were generated as previously described [31]. To normalize for transfection efficiency, the pGL4.54 [luc2/TK] vector (Promega, # E5061) was co-transfected as an internal control. The ACE-tRNA expression constructs—pUC57-4xACE-tRNA^Ser^_UGA_ and pUC57-4xACE-tRNA^Arg^_UGA_—used in this study were previously reported [10]. For all transfection experiments except those involving gene dosage variations of ACE-tRNA, the ACE-tRNA constructs were co-transfected at one-tenth the amount of the COL4A5-Nluc reporter plasmid.

For the split-NanoLuc-based heterotrimer assay, LgBiT-tagged nonsense mutant COL4A5 constructs were created by site-directed mutagenesis using the pFC34K LgBiT TK-Neo Human COL4A5: WT plasmid as a template [32]. Mutagenesis was performed with the following primers: S36X_forward: ctatgggtgttctccaggatGaaagtgtgactgcagtggc, S36X_reverse: gccactgcagtcacactttCatcctggagaacacccatag, S1632X_forward: cctgcttggaagagtttcgttGagctcccttcatcgaatg, S1632X_reverse: cattcgatgaagggagctCaacgaaactcttccaagcagg, R1563X_forward: gcatccagccattcattagtTgatgtgcagtatgtgaagctcc, R1563X_reverse: ggagcttcacatactgcacatcAactaatgaatggctggatgc, R1683X_forward: ggacacgaattagcTgatgtcaagtgtgcatg, R1683X_reverse: catgcacacttgacatcAgctaattcgtgtcc.

To generate NMD-sensitive COL4A5 translational reporter constructs, the Luc2 gene in the pKC-4.04 vector (Addgene, # 112085) [33] was replaced with the COL4A5-Nluc gene. Both the pKC-4.04 backbone and the COL4A5-Nluc insert were amplified using PrimeSTAR Max DNA polymerase (Takara, # R045A), with the following primers: pKC-4.04_forward: gcgaattcatcgatagatctgatatc, pKC-4.04_reverse: ggttaattctgacggttcactaaac, COL4A5-Nluc_forward: gtgaaccgtcagaattaaccATGAAACTGCGTGGAGTCAGCCTGG, COL4A5-Nluc_reverse: agatctatcgatgaattcgccgccagaatgcgttcgcacagccgc.

PCR amplicons were purified using the QIAquick PCR purification kit (Qiagen, # 28104), and then assembled using NEBuilder HiFi DNA Assembly Master Mix (NEB, # E2621S). The integrity of the final constructs was confirmed by Sanger sequencing. Splicing of the HBB intron in the NMD-sensitive constructs was assessed by RT-PCR.

### RNA extraction and quantitative RT-PCR analysis

RNA was isolated from transfected cells using TRIzol reagent (Invitrogen, # 15596018) following the manufacturer’s instructions. Briefly, culture supernatants were removed, and cells were washed once with PBS before lysis in TRIzol. One-fifth volume of chloroform was added to the lysate, and the samples were centrifuged at 12,000 × *g* for 10 minutes. The aqueous phase was collected, followed by the addition of an equal volume of chloroform. After a second centrifugation (12,000 × *g*, 10 minutes), the aqueous phase was again collected. RNA was precipitated by adding an equal volume of isopropanol, followed by two washes with 70% ethanol. The RNA pellet was dissolved in TE buffer. RNA concentration and purity were determined using a NanoDrop 2000c spectrophotometer (Thermo Scientific). Purified RNA was reverse transcribed into cDNA using PrimeScript RT Master Mix (Takara, # RR036A) and used for PCR analysis with the following primers: pKC404 HBB FW1: ggctcacctggacaacctc, pKC404 HBB RV1: ccagcacaatcacgatcatattgc. The expected amplicon size was 773 bp for the unspliced transcript and 119 bp for the spliced transcript. COL4A5-Nluc-HBB mRNA expression was analyzed by quantitative reverse transcription PCR using FastSYBR Green Master Mix (Applied Biosystems, # 4385612) on a QuantStudio 6 Flex Real-Time PCR System (Applied Biosystems), following the manufacturer’s instructions.

### Luciferase assays

Both NanoLuc- and split–NanoLuc–based luciferase assays using the Nano-Glo® Dual-Luciferase® Reporter (NanoDLR™) Assay were performed as previously described [31]. Briefly, for the NanoLuc readthrough assay, supernatants were removed from transfected cells cultured in 96-well white-bottom plates. Cells were then lysed with firefly luciferase substrate buffer, incubated for 10 minutes at room temperature, and luminescence was measured.

Subsequently, NanoLuc luciferase substrate buffer was added to each well, followed by a 10-minute incubation at room temperature, and luminescence was again measured. For the split– NanoLuc–based heterotrimer formation assay, both supernatants and cells were used, and luminescence was measured in the same manner as described for the NanoLuc luciferase assay. NanoLuc luminescence was normalized to firefly luminescence. Luminescence was measured using the GloMax-Multi Detection System (Promega).

### Statistics

Statistical parameters are provided in the figure legends. Luciferase assays were conducted in quadruplicate cell cultures. Statistical significance was assessed using Student’s t-test. Differences with P values less than 0.05 were considered statistically significant.

## Results

### ACE-tRNAs induce full-length protein expression in nonsense mutant COL4A5 cDNA-expressing cells

To assess the full-length translation of COL4A5 mRNA, a previously developed NanoLuc-(NLuc-) based reporter system was used [31]. The wild-type COL4A5-NLuc construct encodes a full-length COL4A5 protein fused to a C-terminal NLuc tag, resulting in detectable luminescence upon translation. In contrast, constructs harboring a premature termination codon (PTC) within the COL4A5 coding sequence (S36X (UGA), S1632X (UGA), R1563X (UGA), R1683X (UGA)) lead to truncated translation products that lack the C-terminal NLuc domain, thereby abolishing luminescence (Fig 1A). This system allows for sensitive monitoring of translation efficiency and the effects of nonsense variants.

**Fig 1.**
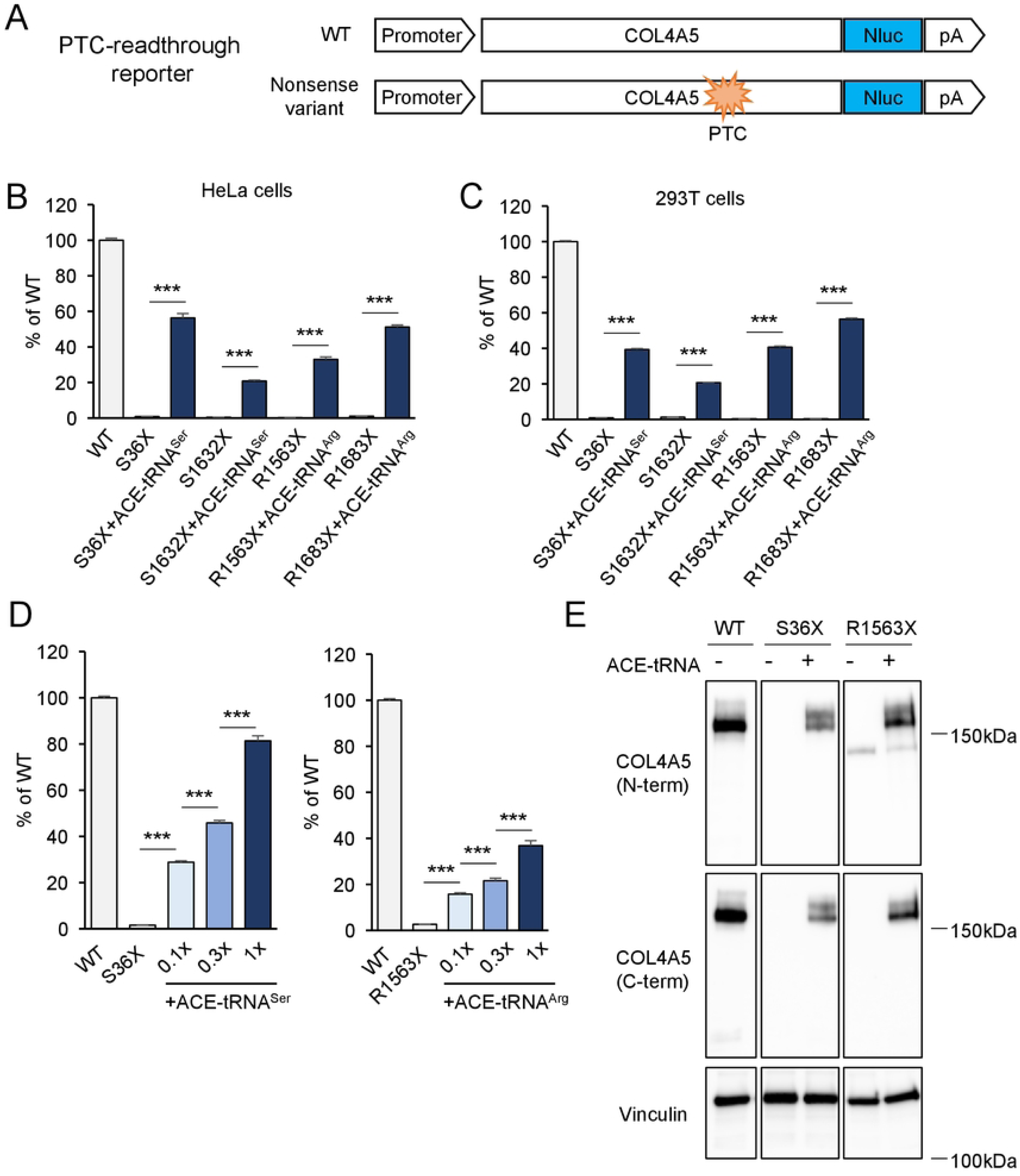
ACE-tRNA-mediated COL4A5 PTC readthrough. (A) NLuc-based COL4A5 translation reporter constructs. The wild-type COL4A5-NLuc construct encodes a full-length COL4A5 protein fused to a C-terminal NLuc tag. Constructs harboring a PTC within the COL4A5 coding sequence lead to truncated translation products that lack the C-terminal NLuc tag. (B, C) Luminescence was measured in lysates from HeLa cells or 293T cells co-transfected with COL4A5-NLuc (WT, S36X, S1632X, R1563X, or R1683X), HSV-TK-Luc2, and either ACE-tRNA^Ser^ or ACE-tRNA^Arg^ constructs. ACE-tRNA treatment promoted translational PTC readthrough in both HeLa and 293T cells, restoring expression of full-length COL4A5 protein. (D) Dose-dependent ACE-tRNA treatment induced readthrough of both COL4A5-S36X and COL4A5-R1563X, with luminescence levels increasing in proportion to the amount of ACE-tRNA plasmid transfected. The ACE-tRNA expression construct was transfected at 0.1×, 0.3×, and 1× of the original plasmid amount, corresponding to the bars shown from left to right in the graph labeled “+ACE-tRNA”. (E) The size of ACE-tRNA-induced full-length COL4A5 protein was assessed by Western blotting. The rescued COL4A5 proteins displayed the expected molecular weight, confirming full-length translation. Vinculin was used as a loading control. Error bars indicate the mean ± SE (n=4). Statistical analysis was performed using Student’s t-test. ***, P <0.005 vs. non-rescued nonsense mutant.

In both HeLa and 293T cells, luminescence was greatly reduced in the four nonsense mutant constructs compared to the wild-type, consistent with premature termination of translation. However, co-expression with ACE-tRNAs resulted in a substantial increase in luminescence across all four mutants (Fig 1B, 1C), indicating restoration of translation through PTC readthrough. The level of luminescence recovery ranged from ∼20% to ∼55%, exceeding that induced by high-dose G418, a conventional readthrough-inducing antibiotic [31]. This readthrough activity was dependent on the ACE-tRNA gene dose (Fig1D). Furthermore, Western blot analysis using both N-terminal and C-terminal anti-COL4A5 antibodies confirmed the presence of full-length COL4A5 protein in ACE-tRNA-treated samples, but not in untreated nonsense mutants (Fig 1E). These results collectively demonstrate that ACE-tRNA effectively facilitates PTC readthrough and restores translation of full-length COL4A5 protein.

### ACE-tRNAs stabilize nonsense mutant COL4A5 mRNA less effectively than an NMD inhibitor but promote PTC readthrough more robustly

Nonsense variants not only prematurely terminate translation but also reduce mRNA levels through nonsense-mediated decay (NMD). This process is regulated by the exon junction complex, which determines whether a transcript is subject to NMD [34]. Consequently, cDNA constructs containing PTCs but lacking introns are typically resistant to NMD. To investigate the effect of ACE-tRNA on mRNA stabilization, NMD-sensitive COL4A5-NLuc constructs were generated by incorporating both HBB exons and introns (Fig 2A). To confirm correct splicing, mRNA was isolated from cells expressing the COL4A5-NLuc-HBB construct and analyzed by RT-PCR using primers spanning two distinct HBB exons. The RT-PCR results demonstrated successful splicing, as the HBB intron was removed (Fig 2B). Introduction of the S36X nonsense mutation led to a reduction in COL4A5-NLuc-HBB mRNA levels (Fig 2C), which was rescued by treatment with the potent NMD inhibitor CC-90009 [35] (Fig 2D). These data showed that the mutant COL4A5-NLuc-HBB constructs were indeed subject to NMD. Co-expression of ACE-tRNA modestly increased S36X COL4A5-NLuc-HBB mRNA levels, but this effect was weaker than that of the NMD inhibitor CC-90009 (Fig 2E). However, the efficiency of PTC readthrough, as indicated by NLuc induction, was substantially higher with ACE-tRNA expression compared to CC-90009 treatment (Fig 2F). While ACE-tRNAs strongly stabilize mRNA in other nonsense mutation contexts [28–30], the degree of mRNA stabilization observed in nonsense COL4A5 overexpressing cells was weaker than expected. Nevertheless, robust PTC readthrough was consistently observed. Together, these findings support previous studies that ACE-tRNAs exert a dual role in suppressing nonsense mutations by promoting both PTC readthrough and mRNA stabilization [13].

**Fig 2.**
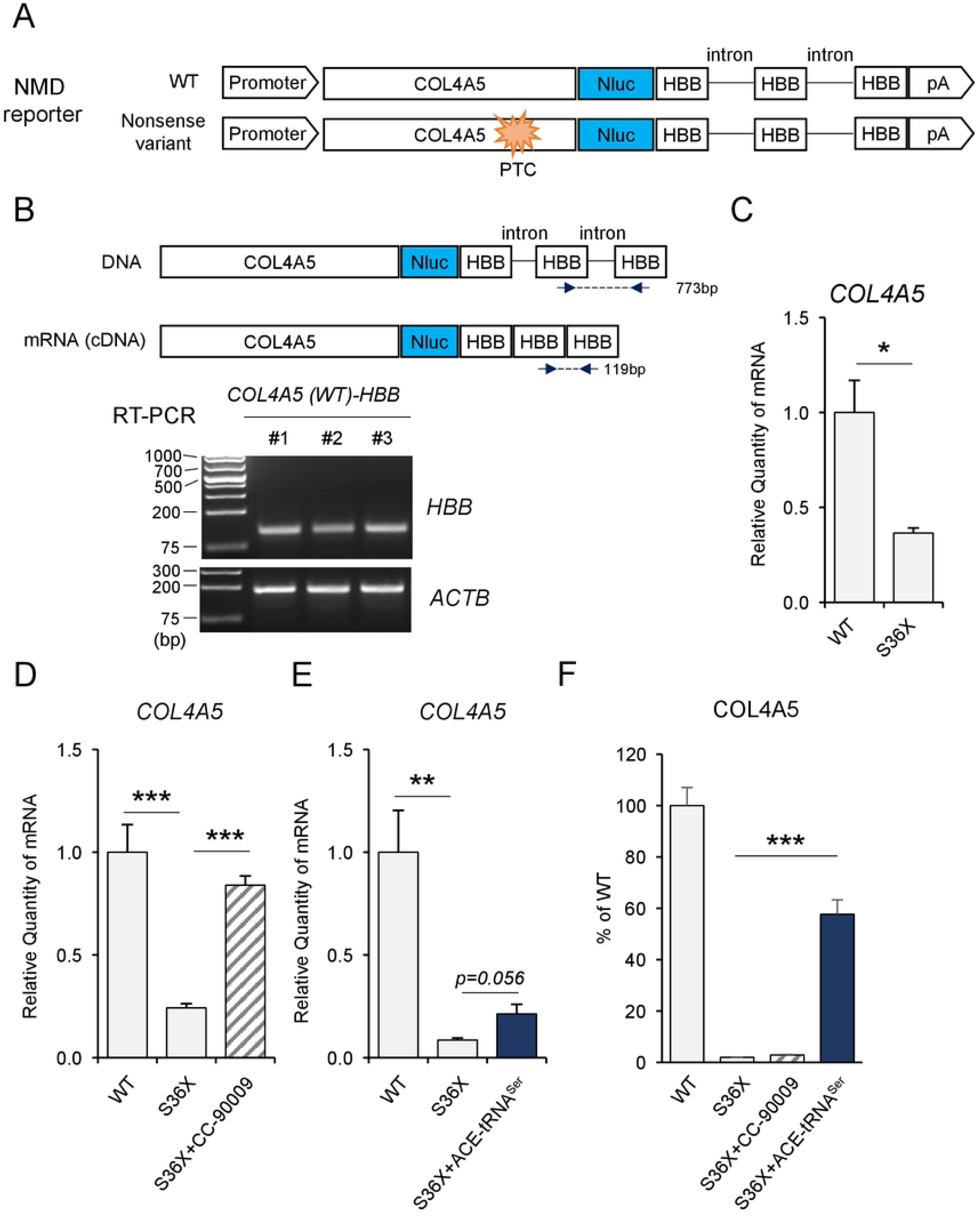
ACE-tRNA stabilizes nonsense-mutant COL4A5 mRNA. (A) Schematic of nonsense-mediated decay (NMD)-sensitive COL4A5-NLuc translation reporter constructs containing introns derived from the human β-globin (HBB) gene. (B) RT-PCR confirmed proper splicing of the COL4A5-NLuc-HBB exon-intron reporter construct. (C) qRT-PCR shows reduced mRNA levels in cells expressing the COL4A5-S36X nonsense variant, consistent with NMD targeting. (D,E) Treatment with the NMD inhibitor CC-90009 or expression of ACE-tRNA^Ser^ partially restored COL4A5-S36X mRNA levels, indicating stabilization via NMD inhibition. (F) PTC readthrough was assessed by measuring NLuc expression following treatment with an NMD inhibitor or ACE-tRNA expression. ACE-tRNA induced PTC readthrough significantly more effectively than NMD inhibition alone, highlighting its dual role in suppressing translation termination and rescuing transcript stability. Error bars indicate the mean ± SE (n=4). Statistical analysis was performed using Student’s t-test. ***, P <0.005, **, P <0.001, *, P <0.05 vs. non-rescued nonsense mutant.

### The full-length COL4A5 protein produced via ACE-tRNA-mediated readthrough is functional

Some of the readthrough products of nonsense variants in COL4A5 mRNA induced by aminoglycosides were non-functional due to the suppression of PTCs by near-cognate tRNAs that insert an incorrect amino acid at the site of the PTC [36]. Since pathogenic missense variants in *COL4A5* are known to impair formation of the α3α4α5 heterotrimers, accurate amino acid incorporation at the PTC site is critical. Thus, restoration of the wild-type residue during PTC readthrough can be as essential as the production of full-length protein for functional rescue. To evaluate the functionality of readthrough products induced by ACE-tRNAs, we employed a split-NLuc-based type IV collagen α3α4α5 heterotrimerization assay that has been previously described (Fig 3A, 3B) [32]. Nonsense mutant COL4A5 constructs showed no luminescence in the split-NLuc heterotrimer assay, indicating a lack of functional trimer formation. In contrast, treatment with the appropriate ACE-tRNAs restored luminescence in these mutants (Fig 3C), with induction levels ranging from ∼9% to ∼45% of wild-type. The relative efficacy across the four mutations mirrored the pattern observed for PTC readthrough efficiency in Fig 1B and 1C. These results suggest that ACE-tRNA not only promotes readthrough of premature stop codons but also facilitates the production of full-length, functionally competent COL4A5 capable of participating in α3α4α5 heterotrimer formation.

**Fig 3.**
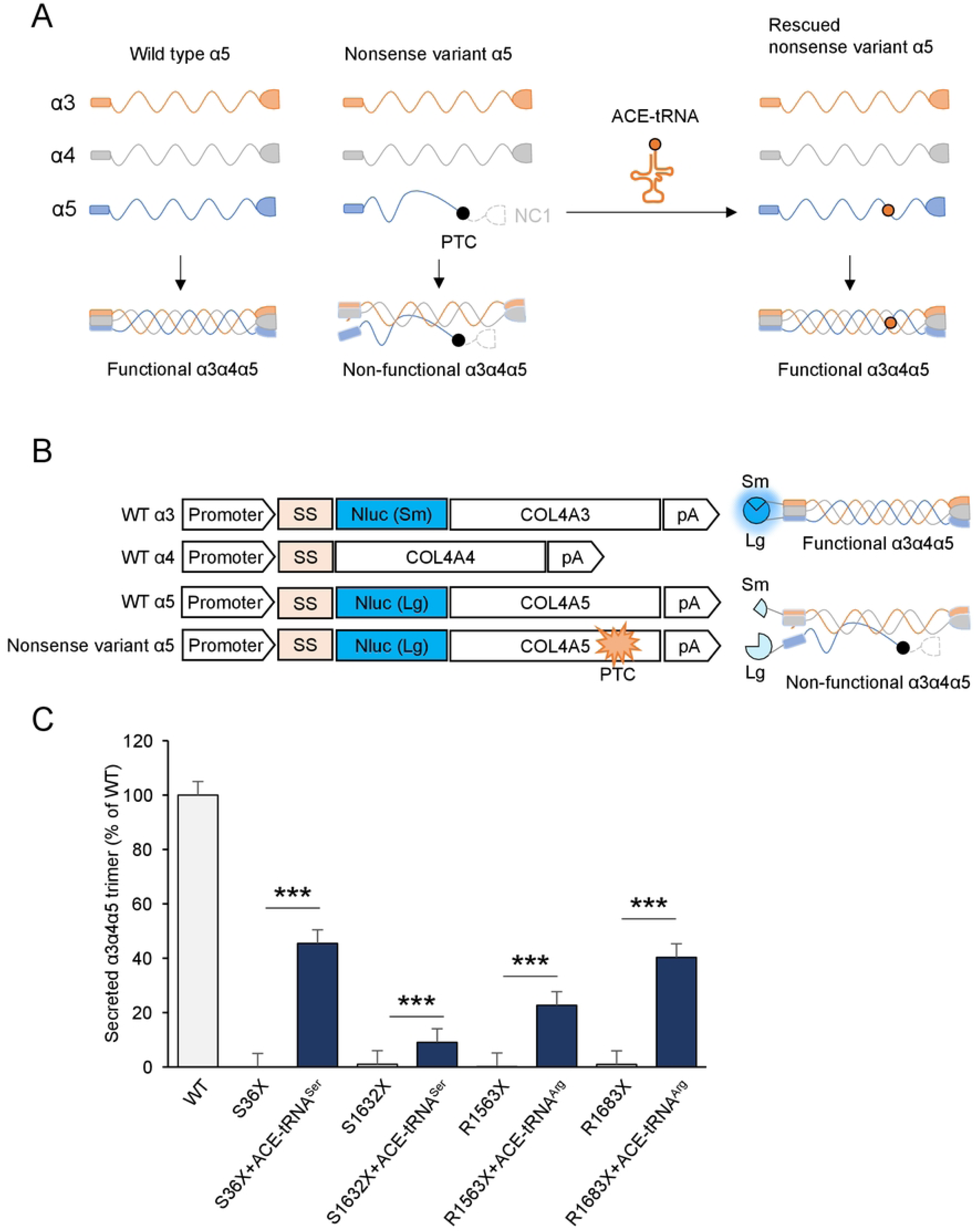
COL4-α3α4α5 heterotrimer formation in ACE-tRNA-mediated PTC readthrough products. (A) Schematic of the split-NLuc-based COL4A3/4/5 heterotrimer assembly reporter system. The split-NLuc fragments (SmBiT and LgBiT) were fused to COL4A3 and COL4A5, respectively, and can generate luminescence only upon proper assembly of the α3α4α5 heterotrimer. Nonsense variant COL4A5 lacks the C-terminal NC1 domain and is unable to form a heterotrimer. (B) Luminescence was measured in culture supernatants from HeLa cells co-transfected with COL4A3-SmBiT, COL4A4, COL4A5-LgBiT, HSV-TK-Luc2, and ACE-tRNA constructs. (C) ACE-tRNA expression restored COL4A5 translation, enabling proper α3α4α5 heterotrimer formation, as evidenced by increased luminescence. These results confirm that ACE-tRNA-mediated readthrough not only restores full-length protein but also supports functional trimer assembly. Error bars indicate the mean ± SE (n=4). Statistical analysis was performed using Student’s t-test. ***, P <0.005 vs. non-rescued nonsense mutant.

## Discussion

Alport syndrome caused by nonsense variants typically presents a more severe phenotype compared to non-truncating mutations, such as missense variants [37, 38]. This is because type IV collagen, the protein affected in Alport syndrome, requires an intact C-terminal domain for proper heterotrimer assembly [39]. Nonsense mutations result in the production of truncated proteins that lack this critical domain, rendering the protein completely non-functional. Therefore, therapeutic strategies aimed at restoring full-length type IV collagen hold significant promise for treating Alport syndrome caused by nonsense variants. In the present study, we demonstrated that ACE-tRNAs effectively induce PTC readthrough, resulting in the restoration of full-length COL4A5 protein expression in cultured cells. Notably, the readthrough efficiency achieved with ACE-tRNA was significantly higher than that observed with aminoglycosides, which are known as classical PTC translational readthrough-inducing drugs (TRIDs) [29]. These findings establish a proof of concept for ACE-tRNA-mediated therapy as a broadly applicable approach for treating nonsense variant-associated Alport syndrome, including cases where conventional readthrough inducers are ineffective.

PTC readthrough activity in mammalian cells was initially discovered using aminoglycoside antibiotics, which were found to suppress premature termination codons and allow translation of full-length proteins [14]. Among these, G418 is recognized as a potent TRID [40] and is widely used as a gold standard in experimental models evaluating the efficacy of PTC suppression. However, despite its strong readthrough activity, G418 is associated with significant nephrotoxicity and ototoxicity, limiting its potential for clinical use [14]. In response to these limitations, gentamicin and other novel classes of PTC readthrough compounds have been developed [41]. As a result, PTC readthrough therapy has emerged as a realistic and promising therapeutic option for genetic disorders caused by nonsense variants. Despite these advances, the efficacy of current PTC TRIDs remains suboptimal and is generally insufficient to achieve a complete cure.

In this context, ACE-tRNAs might represent a significant breakthrough in PTC readthrough therapy. Our findings indicate that ACE-tRNAs mediate significantly greater PTC readthrough efficiency than G418, as demonstrated when comparing the results shown here to those in our previous study conducted under the same experimental conditions [31]. Furthermore, using a split-NLuc collagen IV functional assay, we showed that ACE-tRNA treatment not only induced PTC readthrough but also increased the levels of functional COL4A5 protein. A key advantage of ACE-tRNAs is the ability to incorporate the correct amino acid at the site of the premature stop codon, so called “seamless” rescue of the PTC, ensuring that the resulting protein maintains wild-type sequence and function [10]. This contrasts with traditional PTC TRIDs like aminoglycosides, which rely on near-cognate tRNA recruitment, often resulting in missense amino acid incorporation that can impair protein function [42]. This distinction is particularly important in the context of Alport syndrome, where many pathogenic missense variants—such as glycine substitutions within the collagenous Gly-X-Y repeats—disrupt the structure and function of type IV collagen [43]. In our previous work, we evaluated readthrough products induced by aminoglycosides and found that certain substitutions resulted in non-functional proteins [31], indicating that not all COL4A5 nonsense mutations are amenable to readthrough therapy with traditional agents. In contrast, ACE-tRNAs bypass this limitation by directing incorporation of the correct amino acid and preserving protein function, thus highlighting the therapeutic potential of ACE-tRNAs for a broader range of *COL4A5* nonsense variants, including those not responsive to aminoglycoside-mediated readthrough.

Nonsense-mediated mRNA decay (NMD) plays a key role in the pre-translational regulation of nonsense variant protein synthesis [34]. Although there are exceptions, the majority of mRNAs containing a PTC are recognized and degraded by the NMD pathway, thereby suppressing the production of truncated proteins. Therefore, stabilization of PTC-containing mRNAs through NMD inhibition is regarded as an alternative and complementary approach to augment PTC readthrough efficiency. In this study, we adapted the well-characterized HBB splicing cassette [33] into our COL4A5-NLuc reporter system to investigate the impact of ACE-tRNA on NMD. The mRNA abundance of the COL4A5-S36X-NLuc-HBB construct was significantly reduced, consistent with active NMD, and was effectively rescued by the NMD inhibitor CC-90009 [35]. In contrast, the mRNA-stabilizing effect of ACE-tRNA on the same construct was modest and markedly weaker than that of the NMD inhibitor. Despite its modest impact on mRNA stabilization, ACE-tRNA treatment effectively induced full-length protein expression, whereas the NMD inhibitor did not. This observation is consistent with the mechanism of action of ACE-tRNA, which is designed to promote the incorporation of the correct amino acid at the PTC, thereby enabling the production of full-length functional protein. However, other studies have clearly demonstrated that ACE-tRNA can stabilize mRNA expressed from a gene harboring a genomically encoded nonsense codon [13, 28]. This discrepancy may be attributed to differences in the genes, cell types, or experimental systems used. One limitation of the present study is the use of standard cell lines and an overexpression system, which may not fully recapitulate physiological conditions. Employing more relevant models, such as primary cells derived from nonsense mutant mouse models, could provide deeper insight into the impact of ACE-tRNA on mRNA stabilization. Importantly, our findings also showed that NMD inhibition alone did not lead to full-length protein expression, reinforcing the conclusion that potent PTC readthrough activity, rather than mRNA stabilization, is the critical factor for therapeutic efficacy.

In the context of clinical translation, ACE-tRNAs offer several notable advantages [11]. The compact expression cassette is small enough to be packaged into adeno-associated virus (AAV) vectors, which are widely used in gene therapy [29]. Additionally, because ACE-tRNAs can be delivered as an RNA-based therapeutic modality, they are also compatible with lipid nanoparticle (LNP) delivery systems [30], providing flexibility in therapeutic delivery platforms. Importantly, AAV-based therapies are known for their long-lasting effects, often requiring only a single administration, thereby minimizing the treatment burden. For example, in AAV-based gene therapy for muscular dystrophy, a single dose has demonstrated sustained therapeutic benefits over extended periods [44, 45]. In the case of Alport syndrome, gene replacement therapy remains a challenge because the COL4A5 cDNA is too large to be accommodated within AAV vectors. This limitation underscores the promise of ACE-tRNA as a long-acting therapeutic strategy that directly targets the underlying nonsense variants. By enabling the production of full-length, functional protein, ACE-tRNA represents an ideal approach for directly addressing the genetic defect in Alport syndrome due to nonsense variants.

In summary, the present study proposes the use of ACE-tRNAs as a promising therapeutic strategy for Alport syndrome caused by nonsense variants. ACE-tRNAs exhibit potent PTC readthrough activity and restore the original amino acid, enabling the production of full-length, functional proteins. Importantly, ACE-tRNAs can be customized to target all three types of nonsense codons, each with the appropriate amino acid, offering a precise and adaptable therapeutic platform. Our findings establish a proof of concept for ACE-tRNA–mediated therapy in Alport syndrome and provide a strong rationale for advancing to *in vivo* studies in the near future.

## References

1. Hamosh A, Trapnell BC, Zeitlin PL, Montrose-Rafizadeh C, Rosenstein BJ, Crystal RG, et al. Severe deficiency of cystic fibrosis transmembrane conductance regulator messenger RNA carrying nonsense mutations R553X and W1316X in respiratory epithelial cells of patients with cystic fibrosis. J Clin Invest. 1991;88(6):1880–5. doi: 10.1172/JCI115510. PubMed PMID: 1721624; PubMed Central PMCID: PMCPMC295756.

2. Duan D, Goemans N, Takeda S, Mercuri E, Aartsma-Rus A. Duchenne muscular dystrophy. Nat Rev Dis Primers. 2021;7(1):13. Epub 20210218. doi: 10.1038/s41572-021-00248-3. PubMed PMID: 33602943; PubMed Central PMCID: PMCPMC10557455.

3. Woodley DT, Cogan J, Hou Y, Lyu C, Marinkovich MP, Keene D, et al. Gentamicin induces functional type VII collagen in recessive dystrophic epidermolysis bullosa patients. J Clin Invest. 2017;127(8):3028–38. Epub 20170710. doi: 10.1172/JCI92707. PubMed PMID: 28691931; PubMed Central PMCID: PMCPMC5531396.

4. Lincoln V, Cogan J, Hou Y, Hirsch M, Hao M, Alexeev V, et al. Gentamicin induces LAMB3 nonsense mutation readthrough and restores functional laminin 332 in junctional epidermolysis bullosa. Proc Natl Acad Sci U S A. 2018;115(28):E6536–E45. Epub 20180626. doi: 10.1073/pnas.1803154115. PubMed PMID: 29946029; PubMed Central PMCID: PMCPMC6048497.

5. Kwong A, Cogan J, Hou Y, Antaya R, Hao M, Kim G, et al. Gentamicin Induces Laminin 332 and Improves Wound Healing in Junctional Epidermolysis Bullosa Patients with Nonsense Mutations. Mol Ther. 2020;28(5):1327–38. Epub 20200317. doi: 10.1016/j.ymthe.2020.03.006. PubMed PMID: 32222156; PubMed Central PMCID: PMCPMC7210719.

6. Crawford DK, Alroy I, Sharpe N, Goddeeris MM, Williams G. ELX-02 Generates Protein via Premature Stop Codon Read-Through without Inducing Native Stop Codon Read-Through Proteins. J Pharmacol Exp Ther. 2020;374(2):264–72. Epub 20200506. doi: 10.1124/jpet.120.265595. PubMed PMID: 32376628.

7. Crawford DK, Mullenders J, Pott J, Boj SF, Landskroner-Eiger S, Goddeeris MM. Targeting G542X CFTR nonsense alleles with ELX-02 restores CFTR function in human-derived intestinal organoids. J Cyst Fibros. 2021;20(3):436–42. Epub 20210205. doi: 10.1016/j.jcf.2021.01.009. PubMed PMID: 33558100.

8. Baradaran-Heravi A, Balgi AD, Zimmerman C, Choi K, Shidmoossavee FS, Tan JS, et al. Novel small molecules potentiate premature termination codon readthrough by aminoglycosides. Nucleic Acids Res. 2016;44(14):6583–98. Epub 20160712. doi: 10.1093/nar/gkw638. PubMed PMID: 27407112; PubMed Central PMCID: PMCPMC5001621.

9. Ferguson MW, Gerak CAN, Chow CCT, Rastelli EJ, Elmore KE, Stahl F, et al. The antimalarial drug mefloquine enhances TP53 premature termination codon readthrough by aminoglycoside G418. PLoS One. 2019;14(5):e0216423. Epub 20190523. doi: 10.1371/journal.pone.0216423. PubMed PMID: 31120902; PubMed Central PMCID: PMCPMC6532957.

10. Lueck JD, Yoon JS, Perales-Puchalt A, Mackey AL, Infield DT, Behlke MA, et al. Engineered transfer RNAs for suppression of premature termination codons. Nat Commun. 2019;10(1):822. Epub 20190218. doi: 10.1038/s41467-019-08329-4. PubMed PMID: 30778053; PubMed Central PMCID: PMCPMC6379413.

11. Porter JJ, Heil CS, Lueck JD. Therapeutic promise of engineered nonsense suppressor tRNAs. Wiley Interdiscip Rev RNA. 2021;12(4):e1641. Epub 20210210. doi: 10.1002/wrna.1641. PubMed PMID: 33567469; PubMed Central PMCID: PMCPMC8244042.

12. Dolgin E. tRNA therapeutics burst onto startup scene. Nat Biotechnol. 2022;40(3):283–6. doi: 10.1038/s41587-022-01252-y. PubMed PMID: 35210613.

13. Ko W, Porter JJ, Sipple MT, Edwards KM, Lueck JD. Efficient suppression of endogenous CFTR nonsense mutations using anticodon-engineered transfer RNAs. Mol Ther Nucleic Acids. 2022;28:685–701. Epub 20220504. doi: 10.1016/j.omtn.2022.04.033. PubMed PMID: 35664697; PubMed Central PMCID: PMCPMC9126842.

14. Dabrowski M, Bukowy-Bieryllo Z, Zietkiewicz E. Advances in therapeutic use of a drug-stimulated translational readthrough of premature termination codons. Mol Med. 2018;24(1):25. Epub 20180529. doi: 10.1186/s10020-018-0024-7. PubMed PMID: 30134808; PubMed Central PMCID: PMCPMC6016875.

15. Blanchard SC, Cooperman BS, Wilson DN. Probing translation with small-molecule inhibitors. Chem Biol. 2010;17(6):633–45. doi: 10.1016/j.chembiol.2010.06.003. PubMed PMID: 20609413; PubMed Central PMCID: PMCPMC2914516.

16. Wangen JR, Green R. Stop codon context influences genome-wide stimulation of termination codon readthrough by aminoglycosides. Elife. 2020;9. Epub 20200123. doi: 10.7554/eLife.52611. PubMed PMID: 31971508; PubMed Central PMCID: PMCPMC7089771.

17. Alport AC. Hereditary Familial Congenital Haemorrhagic Nephritis. Br Med J. 1927;1(3454):504-6. doi: 10.1136/bmj.1.3454.504. PubMed PMID: 20773074; PubMed Central PMCID: PMCPMC2454341.

18. Hudson BG, Tryggvason K, Sundaramoorthy M, Neilson EG. Alport’s syndrome, Goodpasture’s syndrome, and type IV collagen. N Engl J Med. 2003;348(25):2543–56. doi: 10.1056/NEJMra022296. PubMed PMID: 12815141.

19. Morrison KE, Germino GG, Reeders ST. Use of the polymerase chain reaction to clone and sequence a cDNA encoding the bovine alpha 3 chain of type IV collagen. J Biol Chem. 1991;266(1):34–9. PubMed PMID: 1985905.

20. Morrison KE, Mariyama M, Yang-Feng TL, Reeders ST. Sequence and localization of a partial cDNA encoding the human alpha 3 chain of type IV collagen. Am J Hum Genet. 1991;49(3):545–54. PubMed PMID: 1882840; PubMed Central PMCID: PMCPMC1683122.

21. Mochizuki T, Lemmink HH, Mariyama M, Antignac C, Gubler MC, Pirson Y, et al. Identification of mutations in the alpha 3(IV) and alpha 4(IV) collagen genes in autosomal recessive Alport syndrome. Nat Genet. 1994;8(1):77–81. doi: 10.1038/ng0994-77. PubMed PMID: 7987396.

22. Barker DF, Hostikka SL, Zhou J, Chow LT, Oliphant AR, Gerken SC, et al. Identification of mutations in the COL4A5 collagen gene in Alport syndrome. Science. 1990;248(4960):1224-7. doi: 10.1126/science.2349482. PubMed PMID: 2349482.

23. Porter JJ, Ko W, Sorensen EG, Lueck JD. Optimization of ACE-tRNAs function in translation for suppression of nonsense mutations. Nucleic Acids Res. 2024;52(22):14112–32. doi: 10.1093/nar/gkae1112. PubMed PMID: 39673265; PubMed Central PMCID: PMCPMC11662937.

24. Pezzini S, Mustaccia A, Aboa P, Faustini G, Branchini A, Pinotti M, et al. Engineered tRNAs efficiently suppress CDKL5 premature termination codons. Sci Rep. 2024;14(1):31791. Epub 20241230. doi: 10.1038/s41598-024-82766-0. PubMed PMID: 39738338; PubMed Central PMCID: PMCPMC11685654.

25. Blomquist VG, Niu J, Choudhury P, Al Saneh A, Colecraft HM, Ahern CA. Transfer RNA-mediated restoration of potassium current and electrical correction in premature termination long-QT syndrome hERG mutants. Mol Ther Nucleic Acids. 2023;34:102032. Epub 20230916. doi: 10.1016/j.omtn.2023.102032. PubMed PMID: 37842167; PubMed Central PMCID: PMCPMC10568093.

26. Awawdeh A, Tapia A, Alshawi SA, Dawodu O, Gaier SA, Specht C, et al. Efficient suppression of premature termination codons with alanine by engineered chimeric pyrrolysine tRNAs. Nucleic Acids Res. 2024;52(22):14244–59. doi: 10.1093/nar/gkae1048. PubMed PMID: 39558163; PubMed Central PMCID: PMCPMC11662663.

27. Specht C, Tapia A, Penrod S, Soriano GA, Awawdeh A, Alshawi SA, et al. An engineered glutamic acid tRNA for efficient suppression of pathogenic nonsense mutations. Nucleic Acids Res. 2025;53(12). doi: 10.1093/nar/gkaf532. PubMed PMID: 40539513; PubMed Central PMCID: PMCPMC12204705.

28. Ko W, Porter JJ, Spelier S, Sorensen EG, Bhatt P, Gabell JT, et al. ACE-tRNAs are a platform technology for suppressing nonsense mutations that cause cystic fibrosis. Nucleic Acids Res. 2025;53(13). doi: 10.1093/nar/gkaf675. PubMed PMID: 40650978; PubMed Central PMCID: PMCPMC12255306.

29. Wang J, Zhang Y, Mendonca CA, Yukselen O, Muneeruddin K, Ren L, et al. AAV-delivered suppressor tRNA overcomes a nonsense mutation in mice. Nature. 2022;604(7905):343–8. Epub 20220323. doi: 10.1038/s41586-022-04533-3. PubMed PMID: 35322228; PubMed Central PMCID: PMCPMC9446716.

30. Albers S, Allen EC, Bharti N, Davyt M, Joshi D, Perez-Garcia CG, et al. Engineered tRNAs suppress nonsense mutations in cells and in vivo. Nature. 2023;618(7966):842–8. Epub 20230531. doi: 10.1038/s41586-023-06133-1. PubMed PMID: 37258671; PubMed Central PMCID: PMCPMC10284701.

31. Omachi K, Kai H, Roberge M, Miner JH. NanoLuc reporters identify COL4A5 nonsense mutations susceptible to drug-induced stop codon readthrough. iScience. 2022;25(3):103891. Epub 20220208. doi: 10.1016/j.isci.2022.103891. PubMed PMID: 35243249; PubMed Central PMCID: PMCPMC8866893.

32. Omachi K, Kamura M, Teramoto K, Kojima H, Yokota T, Kaseda S, et al. A Split-Luciferase-Based Trimer Formation Assay as a High-throughput Screening Platform for Therapeutics in Alport Syndrome. Cell Chem Biol. 2018;25(5):634–43 e4. Epub 20180308. doi: 10.1016/j.chembiol.2018.02.003. PubMed PMID: 29526710.

33. Baird TD, Cheng KC, Chen YC, Buehler E, Martin SE, Inglese J, et al. ICE1 promotes the link between splicing and nonsense-mediated mRNA decay. Elife. 2018;7. Epub 20180312. doi: 10.7554/eLife.33178. PubMed PMID: 29528287; PubMed Central PMCID: PMCPMC5896957.

34. Kurosaki T, Popp MW, Maquat LE. Quality and quantity control of gene expression by nonsense-mediated mRNA decay. Nat Rev Mol Cell Biol. 2019;20(7):406–20. doi: 10.1038/s41580-019-0126-2. PubMed PMID: 30992545; PubMed Central PMCID: PMCPMC6855384.

35. Lee RE, Lewis CA, He L, Bulik-Sullivan EC, Gallant SC, Mascenik TM, et al. Small-molecule eRF3a degraders rescue CFTR nonsense mutations by promoting premature termination codon readthrough. J Clin Invest. 2022;132(18). Epub 20220915. doi: 10.1172/JCI154571. PubMed PMID: 35900863; PubMed Central PMCID: PMCPMC9479597.

36. Roy B, Leszyk JD, Mangus DA, Jacobson A. Nonsense suppression by near-cognate tRNAs employs alternative base pairing at codon positions 1 and 3. Proc Natl Acad Sci U S A. 2015;112(10):3038–43. Epub 20150302. doi: 10.1073/pnas.1424127112. PubMed PMID: 25733896; PubMed Central PMCID: PMCPMC4364220.

37. Bekheirnia MR, Reed B, Gregory MC, McFann K, Shamshirsaz AA, Masoumi A, et al. Genotype-phenotype correlation in X-linked Alport syndrome. J Am Soc Nephrol. 2010;21(5):876–83. Epub 20100408. doi: 10.1681/ASN.2009070784. PubMed PMID: 20378821; PubMed Central PMCID: PMCPMC2865738.

38. Yamamura T, Horinouchi T, Nagano C, Omori T, Sakakibara N, Aoto Y, et al. Genotype-phenotype correlations influence the response to angiotensin-targeting drugs in Japanese patients with male X-linked Alport syndrome. Kidney Int. 2020;98(6):1605–14. Epub 20200724. doi: 10.1016/j.kint.2020.06.038. PubMed PMID: 32712167.

39. Sundaramoorthy M, Meiyappan M, Todd P, Hudson BG. Crystal structure of NC1 domains. Structural basis for type IV collagen assembly in basement membranes. J Biol Chem. 2002;277(34):31142–53. Epub 20020422. doi: 10.1074/jbc.M201740200. PubMed PMID: 11970952.

40. Burke JF, Mogg AE. Suppression of a nonsense mutation in mammalian cells in vivo by the aminoglycoside antibiotics G-418 and paromomycin. Nucleic Acids Res. 1985;13(17):6265–72. doi: 10.1093/nar/13.17.6265. PubMed PMID: 2995924; PubMed Central PMCID: PMCPMC321951.

41. Spelier S, van Doorn EPM, van der Ent CK, Beekman JM, Koppens MAJ. Readthrough compounds for nonsense mutations: bridging the translational gap. Trends Mol Med. 2023;29(4):297–314. Epub 20230222. doi: 10.1016/j.molmed.2023.01.004. PubMed PMID: 36828712.

42. Keeling KM, Xue X, Gunn G, Bedwell DM. Therapeutics based on stop codon readthrough. Annu Rev Genomics Hum Genet. 2014;15:371–94. Epub 20140418. doi: 10.1146/annurev-genom-091212-153527. PubMed PMID: 24773318; PubMed Central PMCID: PMCPMC5304456.

43. Gibson JT, Huang M, Shenelli Croos Dabrera M, Shukla K, Rothe H, Hilbert P, et al. Genotype-phenotype correlations for COL4A3-COL4A5 variants resulting in Gly substitutions in Alport syndrome. Sci Rep. 2022;12(1):2722. Epub 20220217. doi: 10.1038/s41598-022-06525-9. PubMed PMID: 35177655; PubMed Central PMCID: PMCPMC8854626.

44. McKee KK, Yurchenco PD. Amelioration of muscle and nerve pathology of Lama2-related dystrophy by AAV9-laminin-alphaLN linker protein. JCI Insight. 2022;7(13). Epub 20220708. doi: 10.1172/jci.insight.158397. PubMed PMID: 35639486; PubMed Central PMCID: PMCPMC9310540.

45. Birch SM, Lawlor MW, Conlon TJ, Guo LJ, Crudele JM, Hawkins EC, et al. Assessment of systemic AAV-microdystrophin gene therapy in the GRMD model of Duchenne muscular dystrophy. Sci Transl Med. 2023;15(677):eabo1815. Epub 20230104. doi: 10.1126/scitranslmed.abo1815. PubMed PMID: 36599002; PubMed Central PMCID: PMCPMC11107748.

